# Real-Time Human Interaction with Virtual Swarms in Shared Physical Space

**DOI:** 10.1101/2025.06.15.659521

**Authors:** David Mezey, Palina Bartashevich, Ghadi El Hasbani, Pawel Romanczuk, Heiko Hamann, Dominik Deffner, David James

## Abstract

Human-swarm interaction (HSI) explores how humans engage with distributed collective systems, aiming to incorporate human cognition into scalable and robust robotic swarms. While most HSI research focuses on remote teleoperation via engineered interfaces, real-world integration of swarms into everyday tasks requires natural, embodied interactions in shared physical spaces. To address the limitations of traditional teleoperation studies, and the high resource demands of using physical robot swarms for HSI research, we introduce CoBe XR, a spatial augmented reality system that projects virtual swarms into the physical environment of the human operator. CoBe XR enables real-time, fine-grained, natural interaction between humans and swarm-like agents through full-body movement without dedicated control interfaces or prior training. As a proof-of-concept, we present a behavioral study involving 40 participants who influenced swarm behavior solely through walking. Our results show that human participants were able to adapt to the collective dynamics of the swarm and control it through natural perception-motion control in a shared physical space. We argue that similar extended reality systems can not only reveal how humans perceive and adapt to collective dynamics, but they offer a general platform to understand human behavior or an intermediate solution to design embodied robot swarms.

## I. Introduction

Robot swarms are truly scalable systems capable of robust collective behaviors aimed to tackle challenging problems. While within the swarm relatively simple, local behavioral rules between individual robots can lead to complex group-level dynamics, the overarching external control of such distributed systems by human operators is far from trivial ^[1]^, especially with a large number of robots^[2]^. Human-swarm interaction (HSI) is an interdisciplinary field studying how humans interact with collective systems and how this interaction mutually shapes the behavior of the partaking humans and swarms alike^[3]^. HSI aims to harness the complementary strengths of human cognition and autonomous swarm systems^[1]^, specifically combining the high-level reasoning and contextual awareness of human operators with the distributed robustness and scalability of swarms. By developing engineered control interfaces or facilitating natural interactions between operators and swarms^[4]^, a composite, semi-decentralized, human-swarm system arises, that holds significant potential to enhance both robotic capabilities, and human perception and decision-making. Similar human-in-the-loop collective systems have been proposed to address a wide range of real-world challenges, such as search and rescue, surveillance, and fire or hazard mitigation^[5,6]^. A major obstacle to deploying swarm systems in these contexts remains achieving effective and natural human control^[7]^.

Most existing HSI research has limited power to study natural human-swarm interactions, as they focus on remotely controlled robot swarms where human operators are placed in a separate environment distant from the swarm itself ^[8,9]^. One key objective of recent HSI research has been to improve such teleoperation paradigms through the development of augmented intermediate interfaces. These augmented control platforms place the operator in a virtual environment with additional visual cues to support human decision-making during teleoperation^[10–14]^, including aggregated information about the collective dynamics of the swarm^[15–18]^. Although such solutions might partially integrate spatial information from the environment of both the operator and the swarm into a shared interface, these parties remain physically separated.

The practical deployment of human–swarm hybrid systems may require the integration of robot swarms within the operator’s environment to fully realize their potential. As these systems become part of daily life, humans are likely required to naturally coexist with them within the same physical space. In this case, not all users are expected to be trained operators or have access to specialized control interfaces. Instead, in such scenarios, humans have to rely on their innate perceptual and motor abilities to interpret, adapt to, influence and ultimately control complex and dynamic swarm behavior. This type of interaction is fundamentally different from teleoperation. First, it is the interface itself that mediates and filters the possible interactions during teleoperation, rather than innate human capabilities and embodiment. Second, teleoperation usually involves some form of direct or explicit control over the swarm, such as remotely controlling selected robots, whereas natural interactions are often only implicitly shaping group dynamics through sensory information. Lastly, a shared physical space requires continuous interactions between human operator and swarm that are fine-grained, persistent and occur in real-time^[1]^. Despite its importance for developing safe and collaborative systems^[19]^, research on how humans interact with autonomous swarms as part of their environment, rather than as remote operators, remains limited^[20–23]^.

To provide local interactions between robotic swarms and humans in a shared environment, much of the research effort in HSI is dedicated to developing robot perception systems to recognize human presence and actions to enable reactive and adaptive responses^[24–27]^. However, using swarm robot platforms often requires substantial development and maintenance time, as well as additional resources such as technical expertise in robotics, access to multiple robots, and possibly expensive hardware. As a result, with the primary focus on improving these engineered capabilities, the human side of embodied human-swarm interactions often remains secondary. To address these limitations, we propose camera-projected swarms of virtual agents integrated into the human physical environment as an alternative or complement to physical robots in HSI research. Recent advances in camera- and projector-based spatial augmented reality (SAR) systems^[28,29]^ enable dynamic visualizations on real-world surfaces that respond to user behavior in real time, transforming human-environment interaction^[30,31]^. Building on these capabilities, we argue that such interactive systems with projected agents driven by swarm algorithms can serve as an intermediate step for studying embodied human-swarm interactions. In particular, these systems allow researchers to explore fine-grained human behavior in the presence of interacting virtual swarms before deploying actual robotic swarms. Projected agents, being virtual, offer great flexibility for testing different swarm algorithms and adjusting parameters on the fly, without the challenges of robotic implementation and reprogramming. At the same time, these virtual agents still share the same physical space and some of its inherent constraints with human operators. This not only supports more efficient co-design of swarm behaviors aligned with human interaction strategies, but also reduces the temporal, monetary, and technical barriers typically associated with the interdisciplinary nature of HSI research.

In this paper, we present CoBe XR (Collective Behavior eXtended Reality), a SAR system that uses a ground projection to create a real-time, interactive environment, allowing users to engage with dynamic visual content directly integrated into their environment. CoBe XR is equipped with a motion capture system and a set of projectors, connected via a central computer station, allowing the millimetre-precision tracking of marked objects (actors). The system detects actors within a dedicated tracking arena and updates the projected landscape at 45 Hz in response to their position and movement, creating a real-time interaction between the human participant and the projected image. The response is governed by a computer simulation that defines the rules by which the projection changes over time in the presence of detected actors. Using swarm simulations, we can project virtual swarms that share the collective behavior principles with algorithms employed on robotic swarms. This allows us to use CoBe XR to study real-time local interactions between human participants and virtual swarms in a physically shared space, focusing on fine-grained body movements and the emergent collective patterns in the virtual collective without deploying actual robots or implementing specialized control interfaces. Actors can include one or multiple humans, animals, inanimate objects, robots, or a combination of these. This flexibility positions CoBe XR as a general-purpose interactive XR platform applicable not only to HSI research but also to swarm robotics and to exploring real-time animal behavior, including embodied decision-making in general.

As a proof of concept, we present the system’s application to HSI through a case study with 40 human participants, in which we explore whether humans can naturally interact with, control, and adapt to the observed swarm dynamics while sharing the same space, without relying on supporting interfaces, direct control, or any prior knowledge about the swarm behavior or the system itself. The swarm simulation was configured such that virtual agents aimed to remain in a cohesive group during movement while also avoiding the actor, generating unique and composite response patterns. Participants were tasked with interacting with the ground-projected swarm through natural whole-body locomotion to trigger one of these dynamic movement patterns. The target pattern, known as the fountain effect, is sensitive both to the actor’s movement properties, such as speed or direction, as well as to the swarm’s internal state during interaction. As such, its successful reproduction is a highly non-trivial task for which participants must interpret, understand, and learn not only the actual state of the swarm, but its reaction to fine-grained, natural body movement. With our results we demonstrate how real-time, fine-scale tracking combined with highly responsive visual projection can be used to better understand how humans perceive, adapt to, and influence a co-situated swarm through natural perception-motion control.

The paper is organized as follows. In Section 2, we provide an overview of the literature on the use of extended reality systems in HSI research and compare them with our work. Section 3 describes the hardware and software architecture of CoBe XR. Section 4 provides a description of our human behavioral study conducted using CoBe XR, along with the metrics used to quantify human interaction strategies and swarm responses. At last, we provide our main research questions and our planned analysis that will be carried out in later versions of the manuscript.

## II. Related Work

Much of the development in swarm robotics is based on insights from biological swarms and swarm intelligence, which are grounded in principles of emergence, whereby complex collective behaviors arise from simple local interactions among agents without central control^[3]^. To ensure the swarm operates safely and achieves desired outcomes, it is important that human operators can influence the emergence of collective behaviors and modulate their evolution. To form cause-and-effect relationships between their actions and the swarm’s response, humans need to be able to recognize and differentiate the swarm’s behaviors, anticipate their dynamics while remaining aware of their own actions.

Previous research in human perception of swarms has shown that bio-inspired collective motion (akin to bird flocks or fish schools) is easier for humans to perceive than random or unstructured movement^[32]^. Nevertheless, the emergent swarm behaviors remain harder to follow than the motion of biological entities with clear shape and predictable trajectories, such as a walking individual^[33]^. It was demonstrated that humans can differentiate between different swarm behaviors^[34]^, including flocking, dispersion, rendezvous, and swarming^[35]^, displayed on a 2D screen. However, correctly anticipating transition behaviors such as swarm fragmentation has proven to be challenging for a human operator, depending on the swarm’s configuration^[18]^.

Whether humans can learn to understand and anticipate swarm dynamics with appropriate feedback remains largely unexplored, yet it is crucial for operators who need to effectively influence or evaluate swarm behavior. Tabibian et al. ^[36]^ investigated whether humans could select appropriate swarm behaviors to optimize task performance. Participants were asked to predict which of two swarm states, either “rendezvous” or “deploy”, would be more effective for achieving area coverage in a given scenario. Over the course of 50 trials, participants improved their prediction accuracy from 60% to 80%, demonstrating that they were able to learn and refine their understanding of the swarm dynamics. While the study highlights the potential for human operators to learn how their actions affect swarm behavior, it still involves centralized intervention via algorithm selection. In the current paper, we study how humans can influence decentralized swarms without explicit control, enabling emergent dynamics, such as spontaneous structural transitions, through local interactions alone.

Goodrich et al. ^[37]^ studied what types of human influence, without centralized control, are supported by collective behavior that emerges from Couzin’s bio-inspired flocking model^[38]^. They showed that the lead-by-attraction (or lead-by-repulsion) management mode, whereby leader agents are influenced by both the human and other swarm members rather than solely by the human, allows for more effective control of emergent behaviors. This includes structural transitions (e.g., fission-fusion) without changing the parameters of the algorithm. While this was demonstrated via teleoperation, in a physically shared environment a walking human can effectively act as a “leader agent”, continuously influencing the swarm through embodied motion. Indeed, such proximal interactions, where humans and swarms share the same physical space, allow humans to exert a local influence by their physical presence that propagates throughout the swarm, consistent with its distributed dynamics^[3,9]^. Yet, proximal interactions remain largely understudied in HSI, with only a limited number of case studies reported to date^[9,39]^.

Recently developed augmented reality (AR) and mixed reality (MR) interfaces offer a promising means for HSI to create shared environments where humans and robots can interact and collaborate^[40]^. These systems often rely on head-mounted displays, immersing users in hybrid or fully digital spaces^[23,41,42]^. However, such displays can fundamentally alter user kinematics and disrupt vestibular input, leading to sensory mismatches that reduce the sense of presence^[43,44]^. As our perception of the world is deeply connected with the sense of body ownership, which can be distorted in virtual environments, such alterations may influence human interactions and responses^[44,45]^. Given that swarms have been shown to affect a human operator’s emotional state^[46]^ and time perception^[47]^, the combined cognitive and perceptual load that immersive technologies and swarm behavior puts on human operators is unknown.

To this end, CoBe XR, the system proposed in this paper, enables proximal interactions with camera projected virtual swarms in a physically shared space while allowing humans to remain in their natural environment without wearable hardware that alters natural perception, thereby preserving intact visuo-locomotor responses.

Existing large-scale tracking systems for humans or animals typically do not incorporate camera-projected SAR^[48]^, while systems that do are limited to tabletop setups relying on two-handed interaction^[49]^. CoBe XR uniquely combines large-scale tracking with SAR to enable locomotion-based interaction with a co-situated virtual swarm.

Our work contributes to the state-of-the-art in HSI research that studies proximal interactions. In most of such studies, operators interact with robotic swarms exclusively through gestures^[50,51]^, facial expressions^[52]^, or speech^[52]^, typically while remaining stationary. In contrast, this paper studies interaction through natural locomotion in a lead-by-repulsion mode, where the human’s physical displacement (movement) provides continuous input to the physically neighboring swarm agents locally. In our study, the goal for the human participant was to, using only natural perceptual and motor capabilities, learn to anticipate which movement strategies trigger the desired collective behavior through the propagation of local influence. To the best of our knowledge, no prior work has explored how natural human locomotion alone can be used to shape collective dynamics of a swarm in physically shared spaces.

## III. CoBe XR Architecture

CoBe XR is a SAR system that uses ground projection to create a real-time, interactive environment, allowing users to engage with dynamic visual content directly integrated into their environments. In short, CoBe XR detects participants within a dedicated arena and updates the projected landscape in response to their position and movement, creating the effect of real-time interaction. The dynamic projection landscape is governed by an underlying Simulation Module, which defines the rules by which the environment responds to detected participants. The active projected landscape spans approximately 3.8m×3.8m (see Sec. Projection Stack). At each timestep, the system detects marked objects (actors) using a set of infrared (IR) tracking cameras (see Sec. Detection Stack). The resulting 3D coordinates in physical space are then mapped and linearly scaled into a virtual space, based on the maximum possible coordinates in the physical environment and the corresponding bounds in the simulation. In this virtual space, a dedicated Simulation Module updates the environment based on the detected objects’ coordinates, speed, and local interaction rules defined by an underlying computational model (see Sec. Simulation Module). The simulation is rendered as a dynamic visualization of animated fish against a moving water background (see Sec. Visualization). These visuals are then projected onto the arena floor as a light landscape via four projectors (see Sec. Projection Stack). This change is perceived by participants in the arena, completing the full interaction loop — from the user’s perception of the projected environment, to action within the simulated landscape, followed by a dynamic update in response to that action, its rendering and projection, and finally back to human perception.

Each of the four functional components — Detection, Simulation, Visualization, and Projection — operates at its own independent frame rate. The modules communicate via input-output *json* file streams, synchronizing the entire stack in a sequential manner, i.e., each module’s output serves as the input for the next (see Fig. 1D). This design choice ensures that faster processes wait for slower ones, with the slowest module (Visualization Stack) ultimately determining the overall frame rate. Throughout this work, we use the term one “iteration” or “timestep” to refer to a single cycle from human movement to the resulting updated projection. For individual modules, this corresponds to one input read, one computation pass, and one output generation. All modules run on a central computer (Central Node; see Fig. 1D), except for the Detection Stack, which operates on a dedicated machine (Tracking Laptop; see Fig. 1D). Our software stack supports an overall frame rate of 45 Hz, corresponding to a latency of 22.2 ms between user action and the resulting change in the projected image. This enables the system to register fine-grained spatio-temporal patterns in actor movement and to render a responsive visual environment in real time.

**Fig. 1.**
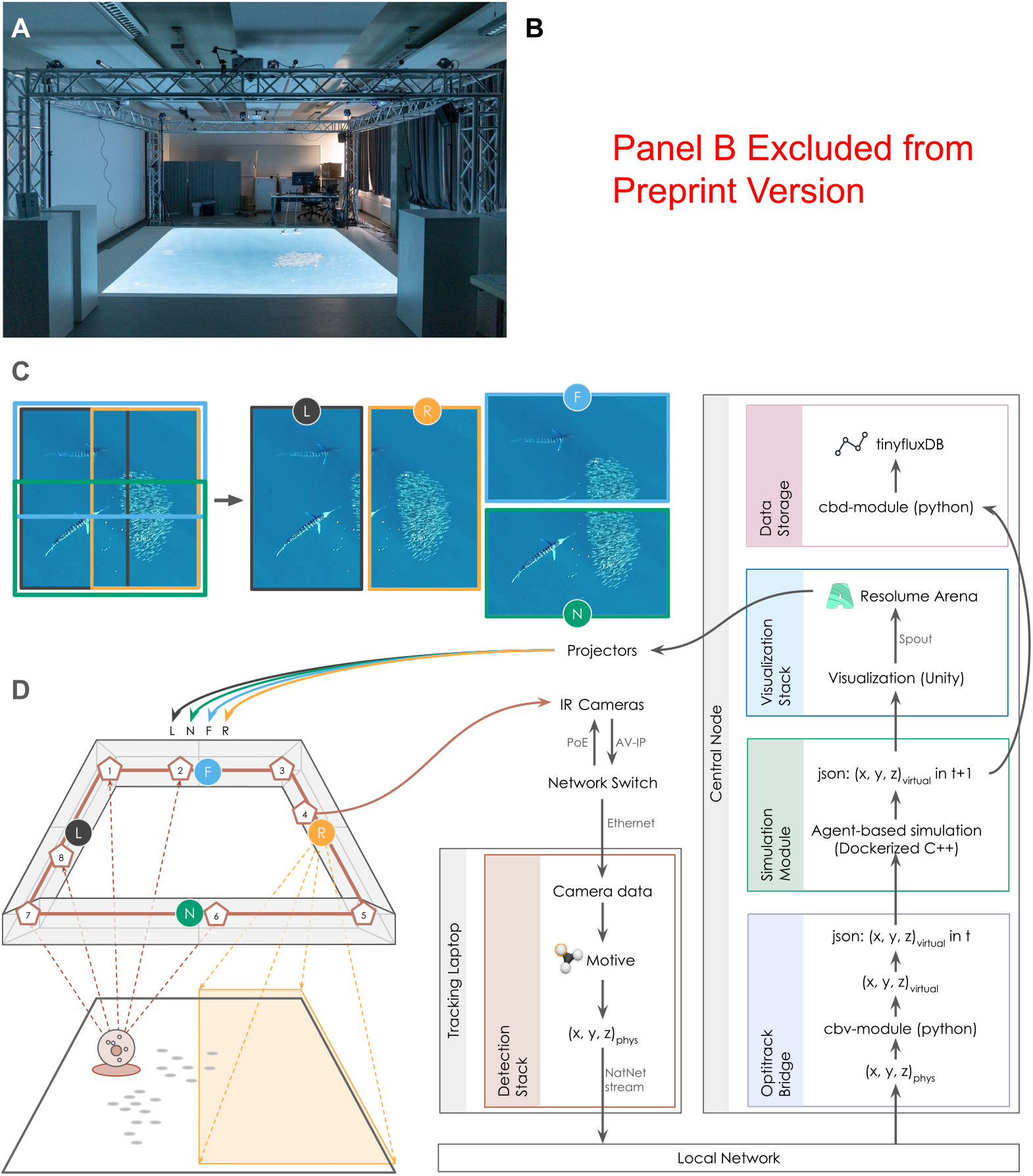
Hardware and Software Architecture of CoBe XR, **A**: Hardware architecture of CoBe XR in operation. **B**: [Excluded from preprint version] A human participant interacting with a swarm of virtual fish in CoBe XR. **C**: Composition of the projected visual landscape. The projected image is sliced into four parts (left, right, near, far) and projected onto the ground by the corresponding projectors (indicated on panel D with matching colors) with partial overlap along the midlines. **D**: Schematic overview of the spatial augmented reality system, showing the connections between hardware components (left) and the software architecture (right). Infrared tracking cameras are indicated with red pentagons, while projectors are denoted with colored circles. The image slice projected by the right projector (R) onto arena ground is shown in orange color. The physical object, equipped with infrared markers, is illustrated as a gray circle containing smaller dark circles. The detected (virtual) position of the tracked object, i.e., actor point, is depicted with a red oval on the ground. The projected fish swarm is schematically shown on the arena ground with light gray ovals. The flowchart illustrates the closed feedback loop in the system, beginning from object detection, passing through the individual components of the software stack, and ending with the projection of the updated landscape back onto the arena. All photographs belong to the authors and the Science of Intelligence Cluster.

### A. Detection Stack

To allow interactions with the system, CoBe XR detects passive infrared markers within the projected arena. These markers reflect infrared light, and they can be placed on any object to be tracked. The reflected infrared light serves as the raw input of the system captured by the eight OptiTrack tracking cameras placed on the four sides and four corners of the surrounding structural metal frame of the arena (see Fig. 1D). The tracking system was recalibrated at the start of each experiment day to assure precise and reliable tracking. Cameras are connected to a dedicated tracking laptop through a network switch (Netgear ProSafe GS728TPP). On the tracking laptop, camera images are registered and the 3D positions of reflective markers are reconstructed within the OptiTrack Motive software^[53]^. This software also allows the definition of ‘rigid bodies’ from unique marker constellations and streams the 3D coordinates of the center of mass of these to the central node through the local network. This way, individual objects can be marked and tracked with a set of attached markers. In this study, we used a 1.5 m cane that human participants could hold in their hands while moving within the system to interact with it. We used a tracking cane instead of direct tracking of the human body to ensure good visibility over the swarm for participants. The cane was tracked via five markers attached to the tip of the cane and allowing for reliable tracking of the cane over time. The detected coordinates together with the IDs of the corresponding rigid bodies are streamed via the local network using Motive with a possible frame rate between 30 and 320Hz. The Central Node reads the streamed coordinates via a dedicated Python module (cbv-module) using NatNet^[54]^ and maps them to a shared virtual coordinate system by scaling them to the absolute size of the virtual space. These new, scaled coordinates are passed to the dockerized Simulation Module through custom *json* files.

### B. Simulation Module

Based on the detected markers’ coordinates from the Detection Stack, the Central Node updates an underlying swarm simulation to emulate an interactive environment according to the movement of the tracked objects. In the current CoBe XR implementation, the simulation module runs a generic agent-based model of schooling prey fish, motivated by predator-prey dynamics, previously validated with empirical observations of schooling sardines under attack by a large marine predator in an open ocean^[55]^. The model computes the positions of virtual prey agents in real time, using simple interaction rules that consider both the neighboring prey agents and the detected markers, with the detected markers interpreted as predator agents that the fish attempt to avoid. The simulation module takes the scaled markers’ coordinates as input together with the virtual prey agents’ coordinates from the previous time step. Interactions between prey agents, as well as those between predator and prey agents, are then calculated according to an underlying computational model (for details, see Sec. III-B1). The virtual prey positions are updated accordingly. Finally, the simulation module generates a custom *json* file with the updated predator and prey coordinates, which serves as input for the visualization stack. The simulation module^[56]^, written in C++, is dockerized for better integration within the otherwise python-based software stack on the central node.

#### 1) Bio-inspired swarm model

Each agent in the model represents an individual “prey” (orange circle in Fig. 2A) moving at a constant speed and changing its orientation according to social forces with the first shell of its nearest neighbors, defined by a Voronoi tessellation^[55]^. A Voronoi tessellation partitions space into cell-like regions, where each region contains all points closest to a given individual^[57]^. Using Voronoi cells provides a biologically plausible way to model information propagation within a group, similar to that during vision-based collective behavior^[58]^. Social forces between agents are modulated by simple behavioral rules such as alignment, repulsion (at short ranges), and attraction (at long ranges). These simple rules give rise to emergent collective behavior of the prey school, such that individuals within the school stay close to each other, and move towards a shared direction with some additive noise. If a predator-agent is in a Voronoi neighborhood subset of an agent, the response to the predator-agent is modulated by the fleeing force when at a certain fleeing distance, such that an agent flees away from the predator towards its rear by turning at a fixed flee angle. The fleeing force is added to the social forces with other prey agents in a Voronoi neighborhood subset of this agent. Due to social interactions, the information about the flee, encoded by the flee force, is propagated from the first shell of the predator’s neighbors to the rest of the prey in the group, resulting in a self-organized collective escape pattern. For more detailed technical implementation, we refer readers to the previous work^[55]^.

**Fig. 2.**
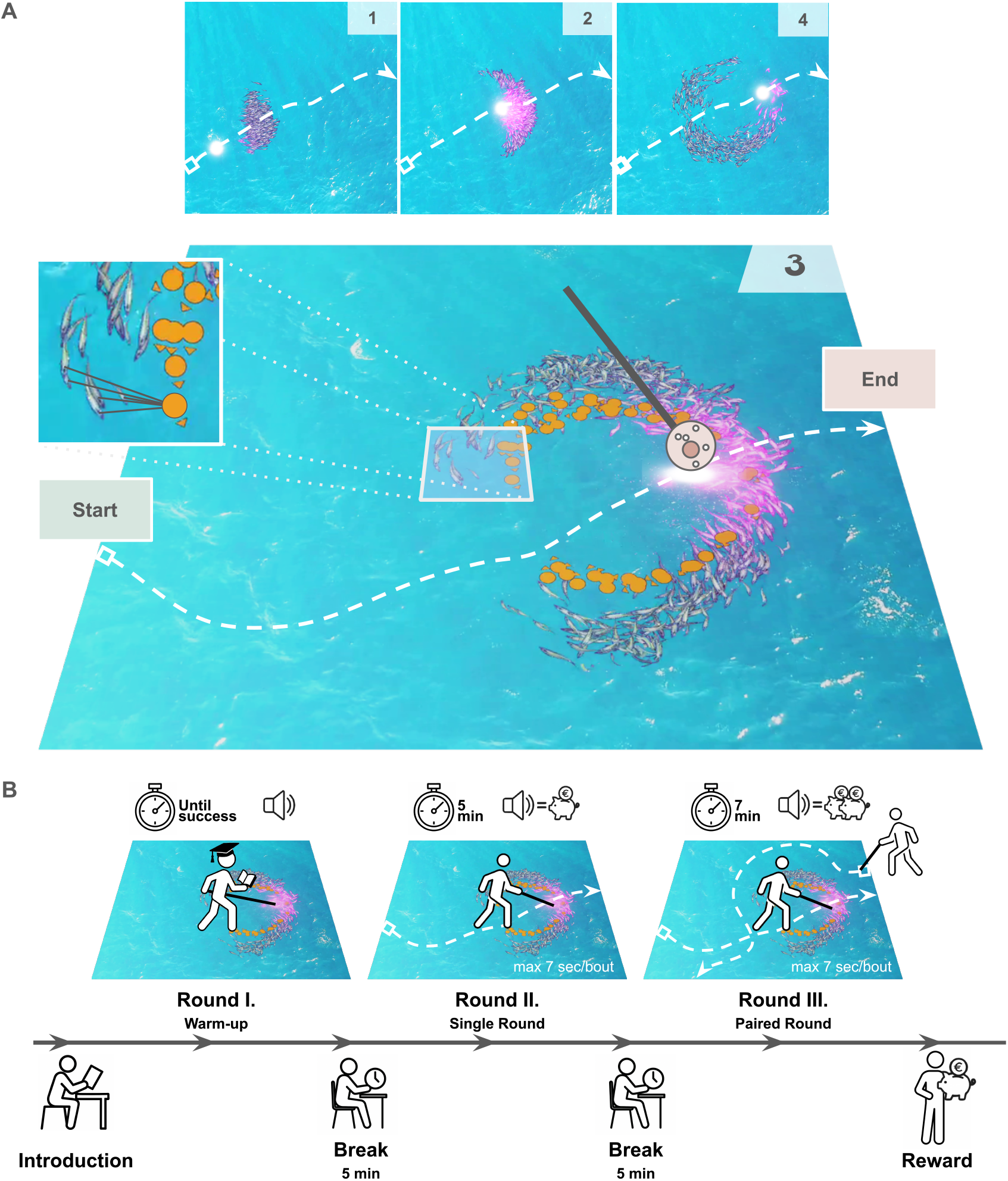
Projected interactive landscape of CoBe XR, illustrating the virtual swarm during the “fountain effect”. **A:** Individual frames of the projected landscape (shown at the top, with frame IDs in the top right corners), captured from the user’s perspective during a single interaction bout with a successfully triggered fountain pattern in the virtual swarm. The third frame, capturing the “fountain effect” is enlarged. On this frame, the trajectory (white dotted line) of a participant is represented by a tracking cane between two sides of the arena (“Start” and “End”). The corresponding predator coordinate (A-point) and virtual fish coordinates (P-Points) in the captured timestamp are shown with a glowing circle and orange dots, respectively. Animated micro swarms appear as smaller animated fish of different color depth and sizes. A single selected P-Point and its corresponding micro swarm is shown in the upper left corner. Note, that during human-swarm interaction only animated fish were visualized without their anchoring P-points. **B:** Timeline illustrating the key stages of a single human–swarm interaction session. After a brief introduction, participants complete a warm-up round, ending with the first successful fountain reproduction. Two experimental rounds follow, separated by 5-minute breaks. Each occurrence of the “fountain effect” was associated with the monetary reward, signaled by sound. In paired rounds, a success by either participant results in a reward for both. If Round III was not feasible, Round II was repeated.

### C. Visualization

The main visualization software is written in C# using the Unity^[59]^ gaming engine on an academic license. At every iteration, the module first reads the new coordinates for each agent and predator from the second most recent *json* file generated by the Simulation Module (for more details see the Supplementary Information). The visualization module then updates the internal state of each game object within Unity using these read positions and corresponding entity IDs. Unity also loads the next frame from the background video file and collates everything into the next finished frame, reflecting the new positions of the virtual agents within the virtual arena. This finished frame is ready to be sent through the Projection Stack. In the current implementation, all detected actor coordinates (A-points) are rendered as large glowing spots, while calculated prey agents’ coordinates (P-points) are rendered as smaller orange dots (see Fig. 2A). Both of these simple visual representations of embodied “predator” and virtual “prey” were colored to be well visible on a blue background, and serve as a simple visualization of the raw output of the Simulation Module.

To facilitate a more dynamic and realistic user experience, we rendered a smaller “microswarm” of five animated fish atop each P-point that is bound to that respective P-point. Each fish in a microswarm varies in size, color and z-distance (plane depth) from the other members, thereby creating the illusion of depth — a seemingly 3-D fish school instead of 2-D representations. These animated fish follow their anchoring P-points according to local interactions based on Voronoitesselation^[60,61]^, partially modified for our purposes (Fig. 2A; for more details see the Supplementary Information). These animated fish both change in color from silver to purple (glow effect) and emit purple particles (sparkle effect) at a rate dictated by an exponential function of the Euclidean distance to the nearest actor, adding another layer of interactive visual effects. After rendering a frame in each iteration of the module, the visualization software sends this image to the Projection Stack via a custom Spout sender^[62]^. The visualization software updates its output at a rate of 50-60Hz depending on the computational load.

### D. Projection Stack

CoBe XR projects the interactive landscape to the arena using four projectors (Optoma 4k 400STx DLP) placed orthogonally on a 2.5 m high static aluminum frame. The beams of the projectors are directed towards the ground using optical mirrors placed directly in front of the projector lenses. Projectors are named by their relative position to the Central Node computer, i.e. left (L), right (R), near (N) and far (F) (Fig. 1D). To minimize shadows resulting from the light occlusion of participants, we project two full, identical copies of the projected visual landscape on top of each other (see Fig. 1C). One of these copies is projected by the L-R projector pair, while the other by the N-F projector pair. Accordingly, each projector is dedicated to visualize approximately one half of the projection landscape. For instance, the L-projector which visualizes a single copy of the left half of the landscape, overlaps with the left half of the N-projector and the left half of the F-projector landscapes. Each of the projected slices are slightly larger (ca. 10%) than a real half of the landscape to allow projection blending with smoothed edges along the midlines of the arena (see Fig. 1C, overlapping rectangles at the midline).

The projectors are orchestrated via Resolume Arena^[63]^. This software allows the transformation of the projected images to (1) align the projected images with the ground surface, (2) to correctly align individual projections with each other, (3) to blend projected images along smoothed borders and the midlines, (4) to correct slight irregularities in the projection surface through local transformations. Furthermore, it establishes a direct connection with the visualization software through a Spout receiver module. In each timestep, the Visualization Module triggers Unity to send the finished image via its Spout sender, which is then received by Resolume Arena’s Spout receiver. This image is then transformed, sliced and distributed to the four projectors according to their respective calibrations in Resolume Arena to produce the final light landscape on the ground. Projectors are updated at a rate of 60Hz.

### E. Data Storage

We extended the software stack with a Data Storage module, which optionally allows the recording of the output of the Simulation Module into a tinyflux^[64]^ database, optimized for time series datasets based on tinyDB^[65]^. This database was populated via Python 3 and was saved as a single *json* file after each experiment. While database writing is active, the corresponding timestamp, the coordinates of all P-points, and each detected actor’s positions scaled into the simulation environment would be saved at each time step. That is to say, the positions of the A-points (i.e., virtual “predators”) corresponding to the physical tracking cane positions in the real arena were saved. To reduce I/O operation overhead, we undersampled the output of the system during database writing, i.e., we saved only every second visualized time frame into the database, yielding an overall ca. 26Hz saving rate. Databases were anonymized by encoding the original timestamps as relative differences from an arbitrary starting point. In this way, no direct link between experiment times and resulting data can be made. We compiled and deposited the resulting anonymized dataset as well as the processed dataset with corresponding metrics and visualizations^[66]^.

## IV. Case Study

As a proof-of-concept for CoBe XR, we conducted an HSI case study that combines real-time, fine-scale tracking with an interactive camera-projected virtual swarm in a shared physical space. Specifically, we tracked the positions of human participants moving within the CoBe XR arena while exerting a repelling leader-based influence^[9,37]^ on the swarm projected onto the arena ground, recording both human and virtual swarm positions for later analysis. While walking in the arena, participants held a tracking cane rigidly in front of their bodies with both hands in a golf-club grip, with the tip of the cane treated as a virtual predator by the underlying swarm simulation. Following the definition of Kolling et al. ^[1]^, this type of human-swarm interaction is *embodied, natural, proximal* and *persistent*, whereby human operators provide a continuous control signal for the swarm using only natural locomotion.

The leader’s influence was created from the underlying agent-based model (see Sec. III-B1) by causing neighboring agents to the virtual predator to be repelled within *R*_*flee*_ = 10. The fleeing direction during repulsion was computed by taking the vector from the predator to the prey agent and rotating it by 30^*°*^ away from the predator’s velocity vector. The angle was chosen based on a previous modeling study, supported by empirically observed escape trajectories in schools of fish attacked by a much faster but less maneuverable predator in the wild^[55]^. Previous work^[55]^ has shown that this flee angle enhances prey escape effectiveness by maximizing individual distance from the predator while enabling quick regrouping to maintain cohesion, resulting in a collective evasion pattern known as the “fountain effect”. The fountain effect can be qualitatively described by (1) a group splitting into two sub-groups in front of a threat, (2) moving around it in arching trajectories that resemble a fountain, and then (3) rejoining into a cohesive, polarized school behind the predator (see Fig. 2A and Fig. 3B1-6).

**Fig. 3.**
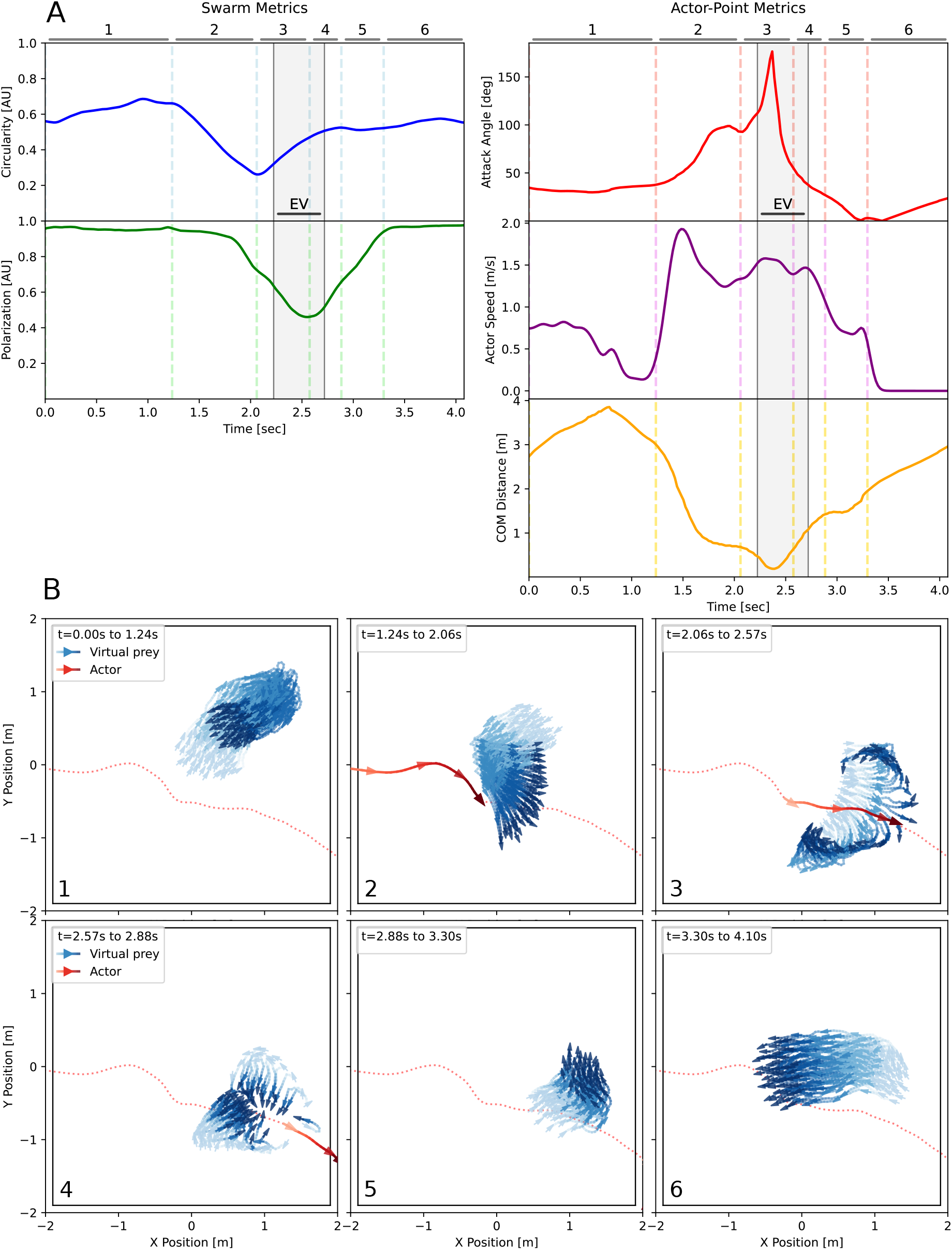
An instance of the recorded interaction bout using CoBe XR, with a successfully triggered fountain pattern in the virtual swarm: **A:** The metrics describing the state of the virtual swarm (on the left) and the movement characteristics of the actor point (on the right) are computed over the duration of the bout. EV denotes the evasion time window detected programmatically (see Sec. Evasion Detection). **B:** Each subfigure shows the temporal evolution of the actor point and swarm agents’ coordinates during the corresponding phase indicated in panel A with matching numbers. A red dotted line traces the full trajectory of the actor across the entire bout, while red and blue arrows represent the actor’s and swarm agents’ movement within each phase, with lighter colors indicating earlier and darker colors later time steps. **1, Resting Phase:** The virtual swarm operates without the influence of the actor, resulting in circular and highly polarized formations. **2, Guiding Phase:** The actor enters the arena and approaches the swarm, which splits in response to the induced repulsion force, resulting in reduced circularity while maintaining high polarization. **3, Arching Phase:** The formed sub-swarms move in arch-like trajectories by either side of the actor, regaining overall swarm circularity but losing polarization; the actor is closest to the swarm’s center of mass and modulates its attack angle. **4, Rejoining Phase:** The swarm regroups behind the actor, increasing circularity while remaining unpolarized. **5, Realigning Phase**: The swarm is in a transient state as it recovers a cohesive, circular, and polarized formation, returning to the resting state **6, Resting Phase**.

As the emergence of the collective escape pattern depends on certain predator’s movement characteristics and attack strategies (e.g., predator speed compared to prey, approach angle to prey), the research question in focus is whether humans can intuitively discover these to reproduce a fountain pattern in a virtual swarm. The task is non-trivial, as participants are unaware about the underlying mechanisms of the swarm dynamics and have to infer the spatio-temporal relationship between their own locomotion (e.g., speed, approach angle, distance to the swarm) and the resulting collective behaviors. This constitutes a form of implicit, decentralized control, where the human has to control the swarm via propagated influence^[9,37]^, i.e., impacting a limited number of agents who in turn influence others leading to a cascading effect through the whole swarm.

To evaluate whether participants could reliably trigger the “fountain effect” in a ground-projected swarm through embodied interaction, we conducted human-swarm interaction experiments with 40 participants, each completing three trials (Fig. 2B, see Methods for details). The first round served as a warm-up, where participants explored the system and attempted to produce a single fountain pattern. Specifically, participants were asked to move with a tracking cane through the swarm in a manner that would cause it to exhibit all three aforementioned qualitative features of the “fountain effect”. The second round, lasting five minutes, aimed to maximize the number of “successful” interactions, i.e., interactions resulting in a fountain effect. For better interpretability in the analysis, participants were encouraged to carry out locomotion-driven interactions in short bouts. Bouts could be initiated from any edge of the arena to any other edge, lasted up to seven seconds, and required forward, uninterrupted locomotion. Each “successful” interaction with a swarm triggered an auditory signal associated with a monetary reward. The third round was either performed solo or in pairs under the same constraint, with paired participants coordinating implicitly, without any communication, to earn shared rewards. To avoid boundary-related distortions, rewards were restricted to “fountains” reproduced inside the arena, closer to its center. In all trials, participants perceived only the animated fish, while the raw swarm simulation output (see Fig.2A, orange dots) was hidden to promote an immersive and engaging interaction experience.

With this experimental design we aim to investigate whether human participants can (1) interact with a projected, autonomously moving fish school in a physically shared space using only locomotion; (2) learn interaction strategies to modify its collective state (i.e., to trigger the “fountain effect”); and (3) adapt these strategies to improve performance in such a task over time. By studying these questions, we evaluate the utility of our SAR system, and similar real-time mixed reality systems, for behavioral research in human-swarm interactions, animal behavior analysis, and swarm robotics

### A. Human-swarm interactions revealed by CoBe XR

In this pilot work, we propose the next analytical steps to answer the aforementioned research questions. We extract individual bouts from the raw recordings (see Sec. Bout Extraction from Continuous Human Movement Recording) and analyze them based on the following five key metrics, which describe both the movement of the virtual swarm and of the human participant’s tracking cane, i.e., the actor point.

The metrics to describe the virtual swarm are (1) circularity, and (2) polarization of the group. We used these metrics primarily to quantify the emergence of the “fountain effect”. The group’s circularity allows us to assess the archness of the escape trajectories on a group level. It defines how round the virtual fish school is, especially during the evasion, taking larger values (between zero and one) for more circular formations. The polarization of the group is calculated as the Euclidean norm of the average normalized velocity vectors of individual prey. If agents move towards a common direction, the group is highly polarized with a polarization value close to one. The polarization drops when the group undergoes fission (split) or fusion (reunion) events, hallmark features of the fountain pattern. The metrics to describe human interaction strategies are (3) the attack angle, (4) the absolute speed of the actor point, and (5) the distance between the actor point and the group’s center of mass (COM). The attack angle is defined by the closed angle between the actor’s velocity vector and that of the prey school’s COM. For detailed definitions of the metrics, see Sec. Methods and Materials.

While the swarm model (Sec. Bio-inspired swarm model) employs the mechanisms that give rise to the fountain-like evasions^[55]^, the structure of the emergent collective pattern is modulated by the trajectory and timing of the repulsive actor’s input. As a result, the model produces a continuum of swarm behaviors (driven by the same mechanism) rather than a single, fixed collective response to an actor. Within this continuum, collective responses range from elongated, chain-like formations to expansive configurations corresponding to the “fountain effect” in its descriptive qualitative form (i.e., split-arches-reunion), our focal pattern of interest.

We identified that a co-occurring steady increase in swarm circularity (above a certain threshold *C*_*L*_), combined with a steady decrease in polarization (below a certain value *P*_*L*_), serves as a strong indicator of fountain-like structures. Specifically, we applied linear regression to identify these trends in both metrics during the respective evasion windows (for details, see Sec. Evasion Detection). Determining the optimal upper and lower threshold values for reliable detection of fountain-like swarm responses remains to be done. We also plan to compare manually assigned fountain scores (see Supplementary Information), a widely used approach for classifying behavioral patterns in field biology^[67]^, with our proposed automatic, metric-based detection method. Manual labeling of the data, at this stage, is yet to be carried out.

Fig. 3 shows an illustrative example of one of the recorded bout instances using CoBe XR, where the actor was able to successfully trigger the “fountain effect” in the virtual swarm. The fine-grained recording of the actor-point’s trajectory, alongside swarm agents’ position data, allowed us to quantify both the actor’s interaction behavior and the swarm’s response in relation to each other (Fig. 3A) to derive potential causal relationships. As the bout started and the actor-point approached the virtual fish school, the group split into two subgroups in front of the virtual predator, elongating its shape and causing a drastic decrease in the group’s circularity (Fig. 3B-2, guiding phase). After the initial split of the group, the resulting subgroups traced arc-shaped trajectories around the actor-point (Fig. 3B-3, arching phase). As a result, swarm circularity increased while polarization decreased, as the formed subgroups moved into the opposite directions (left and right sides) relative to the actor-point. As the actor-point exited the arena, the swarm recovered from the actor’s induced influence. It rejoined into a single group (Fig. 3B-4), recovering circularity to a value comparable with the initial state of the swarm. Finally, virtual agents realigned with each other and the swarm entered a resting state (Fig. 3B-5,6).

To address the research questions posed in this study, we propose the following analysis based on the sort of data as exemplified in Fig. 3. To answer the question if human participants could effectively interact with the projected, autonomously moving fish school in shared space using only locomotion, we first (1) extract successful bouts and quantify the proportion of participants that were able to trigger fountain-like patterns, together with success frequency, and also evaluate the quantitative consistency of the reproduced patterns based on the swarm metrics. To explore learning effects in the participants, we plan to (2) compare individual results over time (intra-individual variability) by analyzing actor metrics during evasion phases. Here, we will examine whether significant changes occur over time and, when applicable, across individual trials. Furthermore, we will (3) compare individual interaction strategies across the participants (inter-individual variability) and explore how specific strategies correlate with success.

Overall, the outlined analytical steps will shed some light on how human participants provide persistent proximal control over a co-located synthetic swarm, if applicable, and how these strategies are learned. With this, we pave the way for a more natural way of interacting with autonomous swarms, exclusively via natural locomotion and posture, relying only on innate human perceptual and embodied motor abilities. Rather than viewing the human as an overarching controller, we highlight interaction with distributed collective systems as an embodied process that includes both influencing swarm behavior and cooperating that do not rely on direct control.

## Acknowledgment

This study was funded by the Deutsche Forschungsgemeinschaft (DFG, German Research Foundation) under Germany’s Excellence Strategy – EXC 2002/1 “Science of Intelligence” – project number 390523135. The funder played no role in study design, data collection, analysis and interpretation of data, or the writing of this manuscript.

We are deeply grateful to Maria Ott, Michael Brück, Mathis Kaiser, Kathleen Waak, and the fantastic team at the Science of Intelligence Cluster for their trust and support in developing CoBe XR, and contribution to the funding of our human behavioral study. We are thankful for TUBS GmbH for co-designing and producing the hardware scaffolding of CoBe XR.

## Contribution Statement

CoBe XR was conceived by DM, PB, and DJ. The system was realized collaboratively: DM designed the overall software architecture and hardware plan, specifically the detection system, PB implemented and integrated the swarm simulations, and DJ developed the visualization software, projection mapping, and co-designed the hardware scaffolding around the arena. Human behavioral experiments were designed by DM and PB with input from DD, and conducted by DM and PB with assistance from GEH, following ethical approval of the ethics committee of the Albrecht Daniel Thaer-Institute of the Faculty of Life Sciences of the Humboldt University of Berlin. The first manuscript draft was written by DM and PB and all co-authors contributed substantially to later revisions. Related work was reviewed by PB with support from DM and HH. Figures were created by DM with feedback from co-authors. Data preprocessing, such as bout separation, implementation of various metrics, and final dataset organization, was performed by GEH under the supervision of DM and PB.

## Data and Code Availability Statement

All relevant code and datasets used for this preprint has been deposited under the following sources: See 68 for metric calculation and bout extraction code, and 66 for the raw, and processed datasets, together with individual bout visualizations. The simulation module is available under 56. For the visualization stack we refer to 69. The general software of CoBeXR is uploaded under 70.

## V. Methods and Materials

### A. HSI Experiments

#### 1) Overview of Human Behavioral Experiments

The HSI experiments consisted of three rounds per participant, with a total of 40 participants overall. Each participant was instructed to perform a series of bouts while being tracked by the system using a handheld tracking cane. The participants’ goal was to reproduce the fountain effect in the virtual fish school as many times as possible, with each successful bout yielding monetary reward.

The cane was positioned in front of each participant and they were instructed not to move the cane independently from their bodies. Instead, they were encouraged to hold the cane as a static extension of their bodies. Bouts could be initiated at any arbitrary time and were allowed between any two edges of the arena. Each bout began when the participant entered the arena and ended either upon exiting or when the maximum time limit of seven seconds expired. This duration was chosen based on the nature of the task, the speed of the virtual fish school, and the dimensions of the active projected arena (approximately 3.8 m × 3.8 m). Time was monitored, and participants were informed if a bout exceeded the time limit. Such bouts were excluded from analysis and not rewarded.

Participants were instructed to move only forward during a bout and were not permitted to stop midway, in order to emulate the natural movement of fish in the open ocean which the agent-based model has been designed to replicate. Occasionally, projected fish could become “stuck” near the arena borders if the participant did not exit the arena and the cane remained positioned between the center and the fish school. Participants were warned about this phenomenon, and no reward was given for fountains reproduced near the arena’s borders respectively. As a result, participants were encouraged to recreate fountain effects as close to the center of the arena as possible, and we excluded bouts where possible boundary perturbations were present due to limited space.

#### 2) Experimental Protocol

Participants waited in a supervised room before the experiment, where communication between them was not allowed. Each participant received a written overview of the experiment beforehand. Prior to each round, they were given time to review a written set of instructions and a description of the upcoming round. Rounds were separated with five-minute breaks. Each round was clearly announced and concluded with verbal instructions.

The first round served as a warm-up. Participants could interact with the virtual fish using the cane and were encouraged to produce at least one fountain. A beeping sound signaled successful reproduction of the fountain effect. While a description of the effect was provided, we deliberately refrained from demonstrating it, requiring participants to learn how to generate the fountain effect independently. The round ended when the participant successfully performed their first fountain within a bout. Although the warm-up round was set to five minutes, most participants achieved this goal in less time. There were no rules on the motion for participants in this round, such that they could freely explore the results of their actions with reactions of the virtual fish school.

In the second round, which lasted five minutes, participants were instructed to perform bouts and produce as many “fountains” as possible. Each successful “fountain” reproduced within the arena boundaries triggered the same beeping sound introduced during the warm-up round. Each beep signified a monetary reward.

The third round was conducted either individually or in pairs, depending on the participant’s availability. Paired rounds lasted seven minutes and involved both participants performing bouts in the same arena. Communication — verbal or non-verbal — was prohibited. If a “fountain” was produced, the reward was granted to both participants, creating a cooperative incentive structure. Each participant could initiate their bout independently, making coordination implicit and voluntary. If paired trials were not feasible, the third round was conducted as a repetition of the individual round following a short break after the second. After completing all three rounds, participants were given their accumulated rewards.

During all rounds, except for the warm up, participants had to follow a set of rules that when followed, would structure the recorded dataset into manageable, and clearly identifiable bouts and encourage continuous, dynamic locomotion of the participants over the whole duration of the experiment. These rules are as follows: participants could start and end their bouts at any edge of the active projection arena at any time during a given round. They were instructed to hold the cane in front of them but to not move it independently from their bodies during locomotion. They were not allowed to stop during bouts, and bouts were limited to a seven second maximum, taking the size of the arena and the speed of the fish into consideration. Only fountains that were reproduced close to the center of the arena were rewarded to avoid participants exploiting possible boundary effects.

### B. Metrics and Data Processing

#### 1) Bout Extraction from Continuous Human Movement Recording

The experiments allowed us to record continuous, unperturbed, voluntary movement of human participants mapped into the same virtual space as our simulation, with an average sampling rate of 43.5, Hz. Each round was then segmented into individual bouts, consistent with the regulations on bout validity communicated to participants (see Sec. Experimental Protocol). Bouts that did not comply with these regulations were discarded. Discarded bouts are labeled in the provided dataset with clear reasons for their exclusion.

To segment each round into bouts, the start and end time points of each bout were identified using a threshold based on the distance of the predator-agent from the arena border. This is a reasonable method for extracting bout boundaries, as all bouts were required to start and end at the borders of the arena. A bout is considered to have started when the predator is within a distance of 0.7*W*_1*/*2_ from the arena center, where *W*_1*/*2_ denotes half the arena width. Once the predator exceeds this distance from the center, the bout is considered to have ended. This approach enables efficient detection of intervals between bouts (when participants are at the arena border) and automatically excludes bouts that occur too close to the border, rendering them invalid.

The resulting bouts are first filtered by duration, retaining only those that consist of *T*_min_ = 30 or more observations, corresponding to a minimum duration of approximately 75 ms. This step removes transient jitters or involuntary short cane movements between the arena edge and the area of interest.

The remaining bouts are then filtered using two velocity thresholds, *V*_*L*_ and *V*_*H*_. The lower threshold *V*_*L*_ serves to discard bouts in which the predator stops moving during the bout, which violates experimental instructions. Specifically, bouts where the predator’s velocity falls below *V*_*L*_ = 0.475m/s for more than 15% of the observations are discarded. In contrast, the upper threshold *V*_*H*_ is used to detect bouts affected by noisy tracking, which can result in sudden spikes in the predator’s velocity. Bouts are discarded if the predator’s velocity exceeds *V*_*H*_ = 6.65m/s during active movement (i.e., when the distance from the arena border is greater than 0.2*W*_1*/*2_).

In paired rounds, it is also important to ensure that jitter in the second cane’s position does not interfere with the first participant’s bout. Therefore, a bout is discarded if the second predator exceeds the high velocity threshold while being located at a distance greater than 0.3*W*_1*/*2_ from the prey center of mass.

Bouts that are not filtered out by the above criteria but still exhibit invalid behavior — such as abrupt velocity jumps, mid-arena stopping, or other unrealistic movements not aligned with the protocol — are manually discarded. These cases are also labeled with explicit exclusion reasons in the dataset^[66]^.

Finally, the bout time windows are each extended by an additional, one second buffer at each end to ensure that activity at the bout boundaries is fully captured.

#### 2) Evasion Detection

Bouts were further processed once identified and extracted from the raw recordings. Specifically, we identified the time windows in which human participants initiated evasion in the virtual swarm. This was necessary to be able to provide a targeted analysis with those time-windows that are the most important for any behavioral interaction between the human participants and the virtual fish school.

The start of the evasion represents the point when the first P-point crosses the line perpendicular to the predator’s velocity vector and gets behind it. In other words the evasion starts if the actor is close enough to the virtual fish school, and moves in such a way so as to induce any individual to move behind it while others still remain in front of the actor. Note, that due to the properties of the system, this can only happen during an active interaction between human and swarm. Once all prey are behind the predator, the evasion is considered to have ended. To control for possible jitters in the predator velocity vector (due to rare, but possible detection noise), especially when the predator is stationary at the beginning of the bout, detected evasion starting points are reset if the number of prey individuals behind the predator drops back down to zero before the end of the evasion has been detected.

We excluded evasions where the actor did not cross the projected swarm. This is considered as a fully successful evasion for the swarm, and by the symmetric nature of a fountain pattern, such evasion strategy can not be observed in such cases. To control for such cases, detected evasion bounds are excluded if no prey is found behind the predator on both sides, left and right, of the velocity vector direction at any point during the detected evasion.

