## Supplementary Information for "Real-Time Human Interaction with Virtual Swarms in Shared Physical Space"

### I. Microswarm Modelling

Each P-point calculated according to the biological model exists within a 2-dimensional plane without overlap. While scientifically valuable, the lack of perceived depth to the render reduced participant immersion in both the scientific and non-scientific uses of the system. The challenge then became how to present the underlying P-points to the participants in such a way so as to seem familiar and believable while not straying too far from the underlying model and thereby change the means of interaction with the system.

The obvious solution would be to render a single fish at each P-point position. However, the obvious caveat was that these rendered fish would then also only exist in the 2-dimensional plane and the school would therefore appear sparse. It is from addressing this issue of fish density that the microswarm idea arose. We would instantiate a larger number of fish, randomly positioned within a certain tolerance around each of the P-points, and which would follow their mother P-point's movements exactly. These fish would all exist within the 2-dimensional plane, but one could simulate depth by layering the fish sprites (models) over one another and tuning sprite color (darker simulating deeper).

The implementation of this model was quick, but also immediately showed certain weaknesses:

1) Because each microswarm's movement would rigidly follows its mother P-point's movements, if the number of P-points was too few or the tolerance distance around a P-point too small, the microswarm's unified movements would become blocky and immediately apparent.
2) The P-points do not have consistent directionality across subsequent iterations and therefore jitter, resulting in every microswarm also noticeably jittering.

A more complex implementation would be needed to address these issues. It was decided that a local microswarm interaction model would be implemented to break up the blocky movements of each microswarm, and that each microswarm would need to be decoupled from, but heavily influenced by, its respective mother P-point to address the jitter while staying true to the underlying biological model.

#### A. Framework

The fundamental mathematical model that would be adapted for the microswarm local interactions is that published by Jacques Gautrais & co. (2012) and then further extended for line of sight considerations by Daniel S Calovi & co. (2014). The joint model outlined by these papers is concerned with defining the change in angular velocity $d\omega$ of each fish at every incremental time step $dt$ based on that individual's neighbors (determined by shared Voronoi boundary) and line of sight.

This combined model is expressed as follows:

$$d\omega_i(t) = -\nu \left[ \frac{dt}{\xi} \cdot (\omega_i(t) - \Omega_{ij}\omega_i^*(t)) - \hat{\sigma} dW \right] \quad (1)$$

$$\omega^* = \hat{k}_W \frac{sgn(\phi_{iW})}{\tau_{iW}} + \frac{1}{N_i} \cdot \sum_{j \in V_i} \left( k_P d_{ij} sin(\theta_{ij}) + \hat{k}_V \nu sin(\phi_{ij}) \right) \quad (2)$$

Where $\Omega_{ij} = 1 + cos(\theta_{ij})$ is a factor that considers the line of sight of the fish as introduced in the Calovi paper, and the values for the constants used are as per the Gautrais paper:

$$\xi = 0.024m$$
$$\hat{\sigma} = 28.9m^{-1}s^{-1/2}$$
$$\hat{k}_W = 0.94$$
$$k_P = 0.41m^{-1}s^{-1}$$
$$\hat{k}_V = 2.7m^{-1}$$

And where:

$\nu$ is the speed of the given fish $i$.

$d_W$ is the wall distance for fish $i$ running along the fish's current orientation vector.

$N_i$ is the sum of the number of fish $j \in V_i$ in fish $i$'s Voronoi neighborhood $V_i$.

$\phi_{iW}$ is the angle between fish $i$'s current orientation and the tank wall.

$\tau_{iW}$ is the time it would take fish $i$ to reach the tank wall at its current speed and orientation.

$\phi_{ij}$ is the heading angle between the orientation vector of fish $i$ and the orientation vector of fish $j$.

$d_{ij}$ is the magnitude of the distance vector between fish $i$ and fish $j$.

$\theta_{ij}$ is the angle between the orientation vector of fish $i$ and the distance vector from fish $i$ to fish $j$.

$dW$ is the incremental change in a Wiener process (equivalent to a randomly sampled Gaussian $\sim \mathcal{N}(0, dt)$).

Implementing this model for each microswarm had those respective microswarms behaving very believably, but entirely ignoring their mother P-point. It is from this foundation that modifications and extensions of the following sub sections were developed.

#### B. Mother is Watching

The first alteration to be made was to have the mother P-point, regardless as to the Voronoi configuration of the microswarm, always be included in the set of neighbors for every fish in the local microswarm. In this way, the mother P-point would always hold some influence over the fish in its associated microswarm. Additionally, the viewing angle factor $\Omega_{iM}$ between a mother P-point $M$ and any fish $i$ within its associated microswarm was always maximal at $\Omega_{iM} = 2$. In this way, the mother P-point's influence would never be attenuated regardless as to its angular position with respect to any given fish $i$ in its local microswarm.

The effect of this modified influence was not as impactful as initially thought, and because the mother P-point's movements are independent of the microswarm, the microswarms would still often break away from their mothers. To strengthen the influence, an additional user-specified weighting term $k_{mom}$ was introduced such that the neighbor term of the $\omega^*$ equation changed from:

$$\frac{1}{N_i} \cdot \sum_{j \in V_i} \Big( \Omega_{ij} \cdot (k_P d_{ij} sin(\theta_{ij}) + \hat{k}_V \nu sin(\phi_{ij})) \Big) \, \forall i \in M \tag{3}$$

To instead:

$$\frac{1}{N_i} \cdot \sum_{j \in V_i} \Big( \Omega_{ij} \cdot (k_P d_{ij} sin(\theta_{ij}) + \hat{k}_V \nu sin(\phi_{ij})) \Big) \cdot (1 - k_{mom}) + k_{mom} \cdot 2(k_P d_{iM} sin(\theta_{iM}) + \hat{k}_V \nu sin(\phi_{iM})) \quad \forall i \in M \tag{4}$$

In which the entire standard neighbor term is scaled down to $(1 - k_{mom})$ its normal value, and the remaining space is weighted for the mother P-point with $\Omega_{iM} = 2$. The value of $k_{mom}$ for the current build is $k_{mom} = 0.2$. However, if a fish is beyond its mother P-point's tank wall (see next section), the value of $k_{mom}$ for that fish becomes 1, removing the influence of any neighboring fish.

This improved the issue but did not entirely prevent micro swarms from escaping their mother P-points.

#### C. Moving Tank Wall

The second alteration made was to the tank wall. The foundational model employs a static tank wall along the arena boundary that is identical for all fish. However, given the nature of the moving P-points, a moving tank wall that would be centered upon the mother P-point was instead deemed more appropriate for controlling microswarm escapes.

As such, a user-specified term $r_W$ was introduced that would dictate the radius of the tank wall that would be centered upon and follow each P-point's movement at every frame. For the current build, that term was set to $r_W = 0.04$. Note that $r_W$ is reported in units normalized to the square arena dimensions of the build which are defined by the coordinates $(0, 0)$, $(0, 1)$, $(1, 0)$, $(1, 1)$.

This modification only affects the wall term of the $\omega^*$ equation, namely the variables $\phi_{iW}$ and $\tau_{iW}$. The value of $\phi_{iW}$ then became defined as the angle between the fish's orientation vector and the tangent line to the tank wall drawn at the point where the fish's orientation vector intersects said wall. The value of $\tau_{iW}$ remained unchanged as the time it would take given the current conditions for fish $i$ to hit the tank wall.

Because the tank wall governing its respective microswarm was now modeled as a moving entity, this meant that fish that were inside the tank in one frame could find themselves outside of the tank in the next if they didn't cover sufficient distance in the last frame. They would then become instantly repelled by the outside wall that now lay directly in front of them and jettison themselves away from the rest of the microswarm. The wall term of the foundational model only accounts for fish crossing a wall in front of them, but has no consideration for a wall that is approaching behind them. This case needed to be addressed as fish that left the moving tank could never return due to the repelling force. To address this, if a fish fell out of the tank, the wall term of the $\omega^*$ equation was ignored (set to 0) until such a time that the fish again resided within the tank boundary. The factor $k_{mom}$ for any fish outside of the tank was set to 1 such that the mother P-point would effectively become the only neighbor.

#### D. Orientation Correction

Subject to the weight of the mother P-point's influence on the local interaction model, individual fish or sometimes the entire microswarm would start to deviate from the path of the mother P-point. To enforce pathing similarity, orientation correction checks were put into place, applied to every fish at every frame.

The mechanism of the orientation correction is to speed up the turning rate of fish that are not oriented within a user-specified angular tolerance of their respective mother P-point. Furthermore, if a fish is positioned beyond a maximum user-defined distance threshold of its mother P-point, then regardless as to that fish's orientation, it will be immediately redirected to face its mother.

The values of the tolerance parameters for the current build are the following:

$$dist_{max} = 4$$
$$dist_{corr} = 1$$
$$\theta_{corr} = \pm\pi/3$$
$$k_{turn} = 1.7$$

Where $dist_{max}$ and $dist_{corr}$ are values normalized to the tank wall radius.

*1) Maximum Distance Check:* The first check in the orientation correction block is to evaluate whether a given fish has exceeded the maximum tank wall radii threshold.

If true, no matter the current angular velocity of the fish, that fish is immediately re-oriented to face its mother P-point, and the orientation correction terminates (skipping the tuning check). The pseudo code for this first check is then:

**if** $dist_{iM} > dist_{max}$ **then**
  $\theta_i = \theta_{i \to M}$
**else**
  next
**end if**

Where $\theta_i$ is the orientation vector of fish $i$, and $\theta_{i \to M}$ is the normalized distance vector from fish $i$ to its mother P-point $M$. Note that this is a different operation from simply aligning the two orientations. In this expression, the orientation of fish $i$ is overwritten to look directly at its mother P-point $M$.

If the given fish has not exceeded the maximum tank wall radii threshold, the tuning orientation correction check then begins.

*2) Tuning Check:* The tuning check only proceeds if both the distance between the fish and its mother point $dist_{iM}$ exceeds $dist_{corr}$, and if the relative heading between the fish and the distance vector to its mother $\angle(\theta_i, \theta_{i \to M})$ is outside of the range $[-\theta_{corr}, \theta_{corr}]$.

Given these conditions, the tuning check proceeds to scale the calculated turning angle of the given fish in the current frame by the value of $k_{turn}$ such that the fish turns faster. However, the value of the resultant adjusted turning angle for the fish for that frame is capped to the range $[-\pi/2, \pi/2]$ to prevent over correction. The pseudo code for this check is then:

**if** $dist_{iM} > dist_{corr}$ **and** $(\angle(\theta_i, \theta_{i \to M}) \notin [-\theta_{corr}, \theta_{corr}])$ **then**
  $d\theta_i = d\theta_i \cdot k_{turn}$
  **if** $d\theta_i \notin [-\pi/2, \pi/2]$ **then**
    $d\theta_i = sgn(d\theta_i) \cdot \pi/2$
  **end if**
**end if**

The full algorithm for orientation correction is then as shown in Algorithm 1.

**Algorithm 1** Orientation Correction

**if** $dist_{iM} > dist_{max}$ **then**
  $\theta_i = \theta_{i \to M}$
**else**
  **if** $dist_{iM} > dist_{corr}$ **and** $(\angle(\theta_i, \theta_{i \to M}) \notin [-\theta_{corr}, \theta_{corr}])$ **then**
    $d\theta_i = d\theta_i \cdot k_{turn}$
    **if** $d\theta_i \notin [-\pi/2, \pi/2]$ **then**
      $d\theta_i = sgn(d\theta_i) \cdot \pi/2$
    **end if**
  **end if**
**end if**

#### E. Distance Correction

Hitherto not discussed is the distance traversed by each fish in a microswarm every frame (calculation cycle). At the most basic level, each fish in a microswarm will traverse the same distance (but not direction necessarily) as its mother P-point every frame. As the distance travelled is the controlled metric, the speed of the fish is then a downstream value that is calculated when also considering the period of the calculation cycle. Given the decoupling between microswarms and P-point, a fish with even a slightly different course (orientation) than its mother P-point will diverge from this P-point over repeated frames. This would still be the case even if said fish quickly corrected its orientation to mimic that of its mother, as the distance travelled by each is always identical and said fish would already be behind. That is to say that unless a fish is always identically aligned with the mother P-point of its microswarm, then due to identical traversed distances each frame, it will invariably fall behind. Thus came into being the distance corrections.

Distance corrections are applied in the form of three multiplicative factors:

1) $k_{fish}$: A fish-specific randomized inherent factor on the range [1.0, 1.08] that is selected when a fish is instantiated and never changes for that fish.
2) $k_{random}$ A fish-specific and frame-specific randomized factor that is independently generated every frame on the range [0.88, 1.12].
3) $k_{geo}$: A fish-specific and frame-specific factor that is calculated based on the current geometry of the fish's position and orientation relative to its mother P-point. With the current settings, this value falls on the range of [0.6, 2].

Then for a given mother P-point $M$ and a given fish $i$ within its associated microswarm, these three factors and the distance travelled by the mother P-point $dist_M$ are all multiplied to generate the final distance travelled by the given fish $i$ in that frame:

$$dist_i = dist_M \cdot k_{fish,i} \cdot k_{random,i} \cdot k_{geo,i} \quad (5)$$

Note that the $k_{fish}$ modifier's range does not fall short of the value of 1.0 as a fish that consistently moves slower than its mother will always fall behind and will therefore consistently hover around the tank wall centered on its mother P-point.

*1) Geometrical Modifier $k_{geo}$:* The $k_{geo}$ factor, as previously indicated, is calculated based on the current geometry of the fish's position and orientation relative to its mother P-point. This factor starts at a value of 1.0 and is only altered if a given fish has fallen outside of the moving circular tank wall around the mother point. If this condition is satisfied, the modifier is then calculated as:

$$k_{geo} = k_{dist} \cdot k_{align} \quad (6)$$

Where $k_{dist}$ is a factor that increases with increasing distance of the fish from its mother P-point, and $k_{align}$ is a factor that increases with increasing alignment between the fish and the distance vector to its mother P-point (as $\angle(\theta_i, \theta_{i \to M}) \to 0$). The underlying logic here is that a fish should only increase its speed if both far from, but oriented towards, its mother P-point.

The distance factor is defined as follows:

$$k_{dist} = \frac{r_s}{1 + e^{(-c_s \cdot suppRads)}} \quad (7)$$

Where the right-hand side of the equation is a sigmoid function, $r_s$ is a user-defined maximum speed-up multiplier due to radial distance beyond the tank wall, $c_s$ is also a user-defined factor that defines how quickly the maximum radial speed is attained (a higher value indicates more quickly), and $suppRads$ is the distance from the fish to its mother normalized to the number of **supplementary** tank wall radii $suppRad = dist_{iM}/wallRadius - 1$. Consequently, the $k_{dist}$ modifier is on the range of [1.0, $r_s$].

The alignment factor is defined as follows:

$$k_{align} = \begin{cases} \cos(\angle(\theta_i, \theta_{i \to M})) \\ \cdot(1 - s_b) + s_b, & \text{if} \angle(\theta_i, \theta_{i \to M}) \in [-\pi/2, \pi/2] \\ s_b, & \text{otherwise} \end{cases} \quad (8)$$

Where $s_b$ is the lowest distance a fish will attain due to alignment mismatch as a percentage of normal distance travelled and $\angle(\theta_i, \theta_{i \to M})$ is the angle between the orientation vector of fish $i$ and the distance vector from fish $i$ to its mother P-point $M$. Consequently, $k_{align}$ is on the range [$s_b$, 1.0].

The user-defined values of the current build are:

$$r_s = 2$$
$$c_s = 5$$
$$s_b = 0.6$$

And therefore $k_{geo}$ is on range [0.6, 2.0]. The product of this combination of values is such that a fish will cover more distance if far away from, but aligned towards its mother (effectively closing the gap). However, if the fish is ahead of its mother P-point, it will slow down to prevent widening the gap.

#### *F. Bringing it Together*

In implementing these microswarms, a more full-bodied school was simulated that could, with enough mother P-points and sufficient layering of individuals across microswarms, have the participants be unable to attribute individual fish to a given microswarm, or even recognize that microswarms were present at all. Individual fish instead appeared to belong to the school at large, coincidentally also the goal of the schooling behavior in nature.

The full algorithm, run at every frame, is shown in Algorithm 2, and is further supplemented by Algorithm 3 on calculating the $\omega_i^*$ value.

**Algorithm 2** Micro Swarm Frame Calculation

**for each** $M \in pPoints$ **do**
  Update pPoint M
  Update tank wall
  **for each** $fish_i \in M$ **do**
    Calculate $\omega_i^*$
    Calculate $d\omega_i$ using the time since the last frame $dt$
    Calculate the new angular velocity for the fish $\omega_i$
    Calculate the new orientation vector for the fish $\theta_i$
    Apply orientation correction as per Algorithm 1
    Calculate the new position of fish $i$
    Apply distance correction
    Apply the updates to the fish
  **end for**
**end for**

**Algorithm 3** Calculating $\omega_i^*$ for Fish $i$

**if** $(x, y)_i \notin tank_M$ **then**
  $k_{mom} = 2$
  $\omega_i^* = k_{mom} \cdot 2(k_P d_{iM} sin(\theta_{iM}) + \hat{k}_V \nu sin(\phi_{iM}))$
**else**
  Determine the point of intersection $(x, y)_{i,tank}$ of the fish heading and tank wall
  Calculate the wall term $\hat{k}_W \cdot sgn(\phi_{iW})/\tau_{iW}$
  Calculate the mother point contribution to the neighbor term $k_{mom} \cdot 2(k_P d_{iM} sin(\theta_{iM}) + \hat{k}_V \nu sin(\phi_{iM}))$
  Calculate the Voronoi neigborhood contribution to the neighbor term $1/N_i \cdot (1 - k_{mom}) \cdot \sum_{j \in V_i} \left( \Omega_{ij} \cdot (k_P d_{ij} sin(\theta_{ij}) + \hat{k}_V \nu sin(\phi_{ij})) \right)$
  Set $\omega_i^*$ =wall term + mother term + Voronoi term
**end if**
**return** $\omega_i^*$

### II. SYSTEM ARCHITECTURE

The CoBe system is built upon a Windows desktop computer with the following specifications:

- **SOMETHING SOMETHING**
- **MORE STUFF**

The architecture of the CoBe system is divided into three distinct modules:

1) Input
2) Model
3) Render & Projection

Each is a self-contained system that can operate independent of the other modules, though all three must be running in parallel for the end result to display as intended.

#### *A. Input*

The Input module is that which is concerned with capturing participant positions and communicating them to the Model module. It therefore serves as the bridge from real-space to simulated space, forming the bridge by way of two Orin nano boards, each of which is equipped with a wide-eye camera lens. The code for this module is written entirely in Python.

The Input module operates in a loop when active:

1) The loop begins when the Input module (main computer) requests image data of the arena from the nanos by interfacing with them over the local network.
2) The nanos receive the request and each sends an image back over the network to the main computer.
3) For each captured image, the module then identifies the object(s) of interest via a machine learning model hosted by RoboFlow. The object of interest is that upon which the machine learning model is trained - a black ball for the current CoBe system.
4) The local coordinates of the object of interest are then converted to simulation space coordinates via the transmitting camera's calibration map and are output to a local system folder as a text file.
5) The loop restarts.

Subsequent iterations of the input module do not overwrite the last output text file, but instead follow an incrementing numerical naming convention. The Input module is only concerned with outputting files, it is not responsible for clearing them afterwards. The Input module frequency currently governs the entire system, cycling at a rate of anywhere between 10 to 12 loops per second (10-12Hz).

The calibration map for each camera is built using the cv2.Aruco Python library. Using this package, an image of 100 Aruco codes arranged in a 10x10 grid is generated and sent to the Unity game engine over the network via TCP. This image is then projected onto the floor through the projectors and each camera captures a single picture. These pictures are sent back to the main computer over the network, which upon reception, are scanned for Aruco codes. Due to the regular arrangement of the Aruco codes in world space, the Aruco codes identified by cv2 in the raw pictures can then be mapped back to the codes in the real world, creating the mapping.

#### *B. Model*

The Model module is where Palina Bartashevich's model is applied. As the model is coded in C++ and was originally coded for a Linux-based implementation, it was dockerized in a Linux container for Windows and communicates with the main operating system via Docker's 'Volumes' functionality.

When active, the docker container also runs in a continuous loop:

1) The code first scans the folder where the Input module output files are written to. To prevent IO permissions errors, the second-highest ranked file is always read (rather than the first).
2) Any files of lower rank than the currently opened file are deleted by the model from the Input module's output folder.
3) The positions of the objects of interest, or "predators" for the purposes of the simulation, are then read from the file and run through the model to calculate the new positions of all the fish for the current frame.
4) The position(s) of the predator(s) and the new positions of all the fish are written into a .json format and output to an output folder that is not the same as the Input module folder.
5) The loop restarts

The Model runs very quickly with a rate exceeding 60Hz. With the Input module outputting at a much lower frequency, it is possible and expected that a given output file from the Input module will be read multiple times by the Model module. However, this does not have any bearing on the calculations performed by the Model module, it's just as though the predator remains stationary for all these intermediate calculations.

Similar to its predecessor in the chain of modules, the Model module is only concerned with outputting to its destination folder, but does not concern itself with cleaning up this destination. That task is held by the next reader in the chain. The output .json file also follows an incrementing numerical naming convention, and so subsequent iterations will never overwrite existing files.

#### *C. Render & Projection*

This module (henceforth RP module) is responsible for accepting text files with the positions of all entities in the system (predators and prey alike), collating everything into a single cohesive render, and then parceling this render out between the projectors for a unified mapped floor projection. There are two distinct entities at play in this module:

- The Unity gaming engine was used to create the render, and heavily relied upon the C# programming language to achieve this.
- Resolume Arena is the paid software that was utilized to perform the projection mapping.

Each of the two entities runs independently of the other. A single loop of the Unity entity runs as follows:

1) Scan the output directory of the Model module for new files. To prevent IO permissions errors, the second-highest ranked file is always read (rather than the first).

2) If the second-highest ranked file has not yet been read, read it and delete all files of lower rank from the directory. Otherwise, next iteration.
3) Process the new data and represent it visually in the Unity render.
4) Stream the Unity window by locally redirecting it within the main computer, via Spout, for anyone who's listening.
5) Restart the loop.

Within Unity, a 'Spout Sender' component sits atop the main camera and passively but continuously sends out the camera view to anyone who will listen.

Similarly, Resolume Arena is rigged to accept a Spout input, to be forwarded through the projectors. However, for a unified image, the Resolume output must first be manually mapped using the tooling available in Output $\rightarrow$ Advanced Output. This manual mapping of the projectors to stitch their seams together need only be done once initially, and then only thereafter if the projection areas on the floor shift (and usually only a touch up at that). A single loop of the Resolume entity simply involves receiving the Spout stream from Unity and then sending it through the projectors.
